# Mathematical modelling elucidates core mechanisms underpinning GnRH pulse generation

**DOI:** 10.1101/245548

**Authors:** Margaritis Voliotis, Xiao Feng Li, Ross De Burgh, Geffen Lass, Stafford L Lightman, Kevin T. O’Byrne, Krasimira Tsaneva-Atanasova

**Affiliations:** Department of Mathematics and Living Systems Institute, College of Engineering, Mathematics and Physical Sciences, University of Exeter, Exeter, EX4 4QF, UK.; EPSRC Centre for Predictive Modelling in Healthcare, University of Exeter, Exeter, EX4 4QJ, UK.; Department of Women and Children’s Health, School of Life Course Sciences, King’s College London, London SE1 1UL, UK.; Henry Wellcome Laboratory for Integrative Neuroscience and Endocrinology, University of Bristol, Bristol, BS1 3NY, UK.

## Abstract

Fertility critically depends on the gonadotropin-releasing hormone (GnRH) pulse generator, a neural construct comprised of hypothalamic neurons co-expressing kisspeptin, neurokoinin-B and dynorphin that drives the pulsatile release of GnRH. How this neural network generates and controls the appropriate ultradian frequency essential for gametogenesis and ovulation is unknown. Here, we present a mathematical model of the GnRH pulse generator with theoretical evidence and *in vivo* experimental data showing that robust pulsatile release of luteinizing hormone, a proxy for GnRH, emerges abruptly as we increase the basal activity of the neuronal network using continuous low frequency optogenetic stimulation of the neural construct. Further increases in basal activity markedly increase pulse frequency. Model predictions that such behaviors are concomitant of non-linear positive and negative feedback interactions mediated through neurokinin-B and dynorphin signaling respectively are confirmed neuropharmacologically. Our mathematical model sheds light on the long-elusive GnRH pulse generator offering new horizons for fertility regulation.

## Introduction

The periodic release of gonadotropin-releasing hormone (GnRH) plays a central role in control of mammalian reproduction and is driven by hypothalamic neuronal networks (1). The operation of these networks at a frequency appropriate for the species is critical for the generation of gonadotropin hormone signals (luteinizing hormone, LH; and follicle-stimulating hormone, FSH) by the pituitary gland, which stimulate the gonads and set in motion gametogenesis and ovulation. However, the mechanisms underlying GnRH pulse generation and frequency control remain poorly understood.

Secretion of GnRH by GnRH neurons located in the hypothalamus into the pituitary portal circulation is controlled by upstream hypothalamic signals (1). The neuropeptide kisspeptin has been identified as a key regulator of GnRH secretion as both humans and rodents with inactivating mutations in kisspeptin or its receptor fail to progress through puberty or show normal pulsatile LH secretion (2-4). Within the hypothalamus, two major kisspeptin producing neuronal populations are found in the arcuate nucleus (ARC) and in the preoptical area (5) or the anteroventral periventricular (AVPV)/rostral periventricular (PeN) continuum in rodents (6). Moreover, the invariable association between neuronal activity in the ARC and LH pulses across a range of species from rodents to primates (7) has been suggestive that the ARC is the location of the GnRH pulse generator, and therefore the ARC kisspeptin neurons, also known as KNDy for co-expressing neurokinin B (NKB) and dynorphin (Dyn) alongside kisspeptin (8), constitute the core of the GnRH pulse generator.

Although animal studies have shown that KNDy neurons are critical for the regulation of GnRH secretion, there has been relatively little understanding on the regulatory mechanisms involved in generating and sustaining pulsatile dynamics. Pharmacological modulators of kisspeptin, NKB and Dyn signaling have been extensively used to perturb the system and study the effect on the activity of a hypothalamic neuronal population (using ARC multiunit activity (MUA) volleys, an electrophysiological correlate of GnRH pulse generator activity, as a proxy) (9), as well as on downstream GnRH/LH pulse dynamics (10-13). For example, it has been shown that kisspeptin (Kp-10) administration does not affect MUA volleys in the ovariectomized rat (11), suggesting that kisspeptin is relaying the pulsatile signal to GnRH neurons rather than generating it. On the contrary, administration of NKB or Dyn modulates MUA volley frequency in the ovariectomized goat (13), suggesting a more active role for these neuropeptides in the generation of the pulses. Deciphering, however, the role of NKB has been problematic, and there exist conflicting data showing either an increase or decrease of LH levels in response to administration of a selective NKB receptor (TACR3) agonist (senktide) (10, 12, 14). Recently, a study combining optogenetics, with whole-cell electrophysiology and molecular pharmacology has shed light on the action of the neuropeptides NKB and Dyn in the KNDy network (15), with the key mechanistic finding that NKB functions as an excitatory signal by depolarizing KNDy cells at the post-synaptic site, while co-released Dyn functions pre-synoptically to inhibit NKB release.

Motivated by the experimental findings described above, we developed a mathematical model of the ARC KNDy network. The model predicts that the KNDy population behaves as a relaxation oscillator: autonomously generating and sustaining pulsatile activity similar to the hypothalamic MUA volleys observed *in vivo* (9, 10). Model analysis reveals that *continuous* basal activity within the ARC KNDy population as well as the positive and negative feedback interactions mediated by NKB and Dyn signaling respectively are critical for pulsatility. We tested the model predictions in vivo using optogenetics and showed that low-frequency *continuous* neuronal activation in the ARC KNDy network initiates GnRH/LH pulsatility in female estrous mice and small changes in basal network firing can have a large impact on pulse frequency. Furthermore, we showed that blocking NKB and Dyn signaling alters the behavior of system in response to low-frequency *continuous* neuronal activation in support of our modelling prediction.

## Results

### A coarse-grained model of the ARC KNDy population

We propose a new mathematical model (Fig. 1A) to study the dynamics of the ARC KNDy population. The model is derived from a network description of the system (see SI) and describes the neuronal population using three dynamical variables: 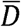, representing the average concentration of Dyn secreted by the population; 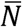, representing the average concentration of NKB secreted by the population; and 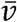, representing the average firing activity of the population, measured in spikes/min. The dynamics of the model variables are governed by the following set of coupled ordinary differential equations (ODEs):

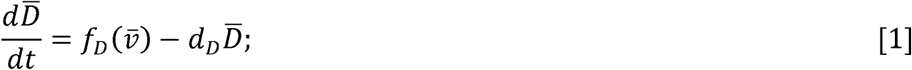

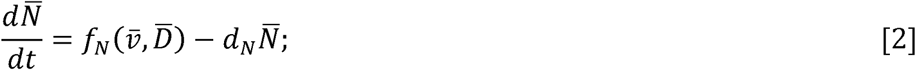

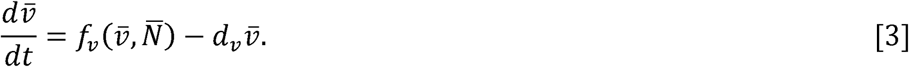

**Figure 1:**
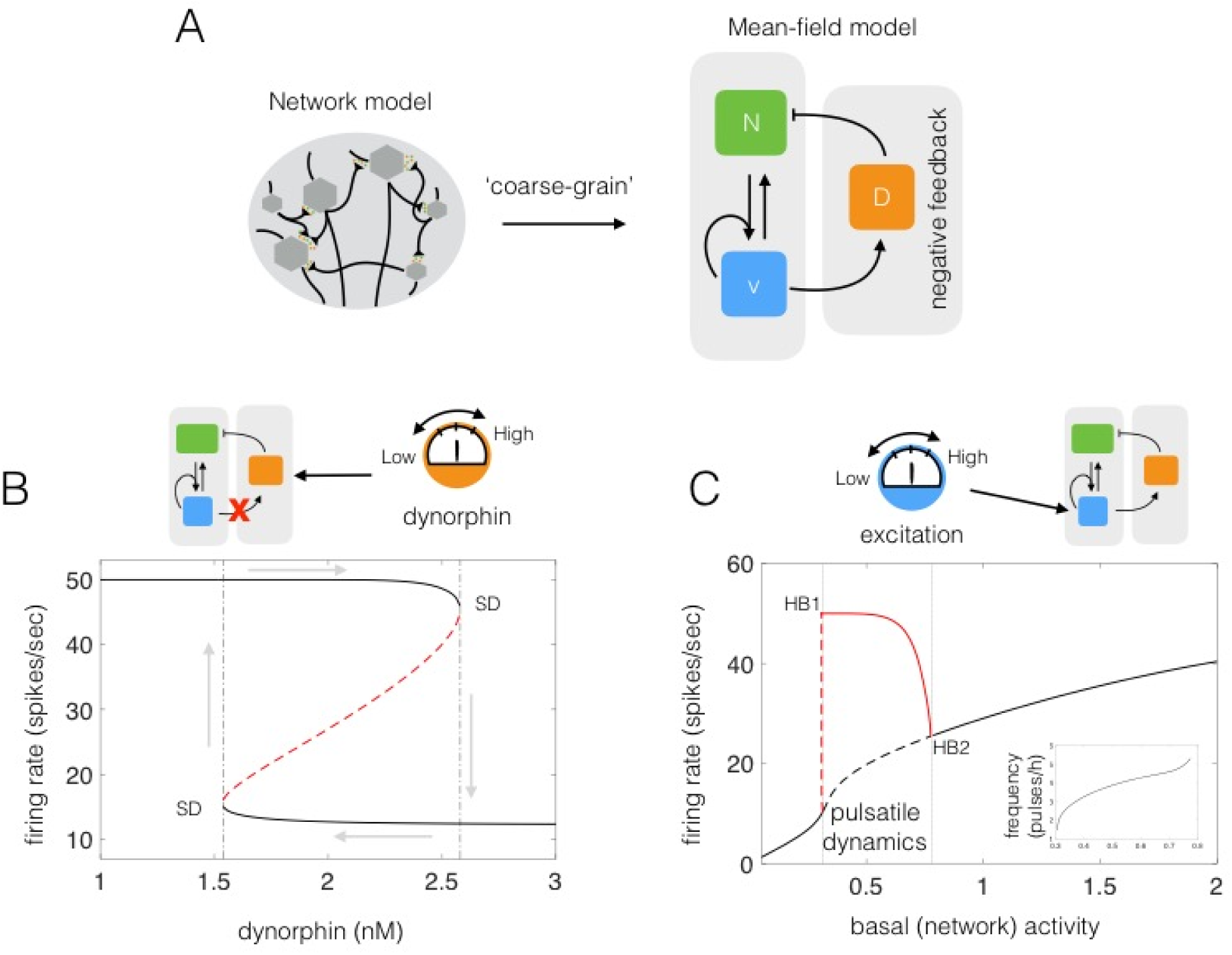
A coarse-grained model gives mechanistic insight into the pulsatile behavior of the ARC KNDy population. (A) We derive a mean-field model of the neuronal population comprising three dynamical variables: 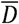, representing the average concentration of Dyn secreted; 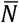, representing the concentration of NKB secreted; and 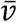, representing the average firing activity of the neuronal population. (B) After disrupting the negative feedback loop (i.e., setting Dynorphin under external control) the system exhibits, for intermediate values of Dynorphin, two stable equilibria (upper and lower solid lines) and an unstable one (dashed red line). At the edges of the bi-stable regime equilibria are lost through a saddle-node bifurcation (SD points). The bi-stability gives rise to hysteresis as the value of Dyn is varied externally (grey arrows). (C) The coarse-grained model predicts how basal neuronal activity affects the system’s dynamics and pulse frequency (inset). As basal activity is increased from zero, high-amplitude, low-frequency pulses emerge after some critical value (HB1 point; Hopf bifurcation). The frequency of pulses continues to increase with basal activity until oscillations disappear (HB2 point; Hopf bifurcation) and the system enters a mono-stable regime (black solid line). The solid red line denotes the amplitude of the pulses. Model parameter values are given in the SI (Tbl. S2).

Parameters *d*_*D*_, *d*_*N*_ and *d*_*v*_ control the characteristic timescale of each variable. In particular, parameters *d*_*D*_ and *d*_*N*_ correspond to the rate at which Dyn and NKB are lost (e.g. due to diffusion or active degradation), while *d*_*v*_ relates to the rate at which neuronal activity resets to its basal level. Functions *f*_*D*_, *f*_*n*_ describe the secretion rate of Dyn and NKB, respectively, while function *f*_*v*_ encodes how the firing rate changes in response to the current levels of NKB and firing rate.

We employ the following sigmoidal (Hill) functions to describe regulatory relationships between the variables. In particular, we set the secretion rate of Dyn and NKB to be:

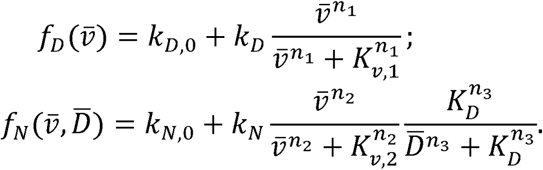

In the equations above, both neuropeptides are constitutively secreted at rates *k*_*D*,0_ and *k*_*N*,0_. Neuronal activity further stimulates secretion of both neuropeptides, and Dyn represses secretion of NKB (15). Since the rate of neuropeptide release is inherently limited by availability of cytoplasmic secretory vesicles at the presynaptic terminals (16), we let secretion rates saturate at *k*_*D*_ and *k*_*N*_, respectively. The effector levels at which saturation occurs are controlled via parameters *K*_*v*,1_, *K*_*v*,2_ and *K*_*D*_. Furthermore, we set:

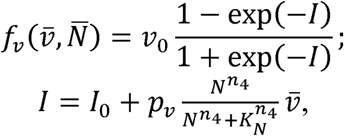

where *v*_0_ is the maximum rate at which the firing rate increases in response to synaptic inputs *I*. The stimulatory effect of NKB (which is secreted at the presynaptic terminal) is mediated via G proteincoupled receptor Tacr3 and is manifested as a short-term depolarization of the postsynaptic neuron (15). In the equation above, we accommodate this effect by letting a synaptic weight that is a sigmoidal function of NKB multiply 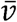. Parameter *K*_*N*_ sets the level of NKB at which its effect is half-maximal, and parameter *p*_*v*_ controls the strength of the synaptic connections between KNDy neurons. Finally, parameter *I*_0_ controls the basal neuronal activity in the population, which could stem from synaptic noise or external inputs. We find that for biophysically relevant parameter values (see SI, Tbl. S1) the model reproduces the synchronized neuronal activity measured from the hypothalamus of ovariectomized rats (10). This finding further supports the hypothesis that KNDy neurons in the ARC constitute the core of the GnRH pulse generator.

### Analysis of the model reveals the KNDy population functions as relaxation oscillator

Having shown that the model can reproduce sustained pulses of neuronal activity (see Fig. 1B), we proceed to investigate the mechanisms driving the phenomenon. We first focus on the role of Dyn-mediated negative feedback using fast – slow analysis (17) of the coarse-grained model (Eqns. [1–3]). Model calibration suggests that Dyn operates at a slower time-scale than NKB (see SI). This time-scale separation, also supported by receptor internalization data (18), allows us to study the dynamics of the fast subsystem, comprised of 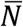 and 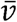, as a function of the slow variable, 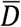, which is treated as a constant (bifurcation) parameter. Our analysis shows that for intermediate values of Dyn the fast subsystem can exist either in a high or a low activity state as demonstrated by the co-existing high and low branches of stable equilibria in Fig. 1B. This bi-stable behavior, stemming from the non-linear, positive feedback between neuronal activity and NKB secretion, leads to sustained oscillations of neuronal activity when combined with slow, Dyn-mediated, negative feedback. In engineering terms, the system behaves as a relaxation oscillator: where the bi-stable subsystem is successively excited (moved from low to high state) by external inputs or noise and silenced (moved from high to low state) as a result of negative feedback. We should note that a relatively slow negative feedback is sufficient for sustaining oscillations, however, the combination of negative feedback with bi-stability is found in many biological oscillators most likely because it confers robustness (19).

Next, to demonstrate the role of basal neuronal activity within the KNDy network in the generation and modulation of oscillatory activity, we treat parameter *I*_0_ as a bifurcation parameter. Our analysis shows that oscillatory behavior is supported within a critical range of *I*_0_ values (see Fig. 1C). As *I*_0_ is increased from zero, high-amplitude, low-frequency pulses emerge via a Hopf bifurcation (Fig. 1C; HB1 point). The frequency of pulses further increases with *I*_0_, until oscillations disappear via a Hopf bifurcation (Fig. 1C; HB point) and the system re-enters a silent (non-oscillatory) regime.

### Continuous optogenetic stimulation of KNDy neurons generates pulsatile LH secretion *in vivo*

To test the model prediction that continuous excitation triggers pulsatile behavior of the system, we stimulated the ARC KNDy population in Kiss-Cre mice (20) using optogenetics. ARC kisspeptin-expressing neurons were transduced with a Cre-dependent adeno-associated virus (AAV9-EF1-dflox-hChR2-(H134R)-mCherry-WPRE-hGH) to express channelrhodopsin (ChR2). AAV-injected, Kiss-Cre mice were implanted with a fiber optic cannula in the ARC and the effects on LH pulsatility of continuous stimulation at different frequencies was tested. After 1 h of controlled blood sampling, low-frequency optic stimulation, 5-ms pulses of blue light (473 nm) at 0.5, 1 or 5 Hz, was initiated and continuously delivered for 90 min. Control mice received no optic stimulation. During the course of the experiment, blood samples (5μl) were collected every 5 min (20). To maximize the effect of optogenetic stimulation, estrous mice were used which display minimum intrinsic pulse generator activity (21). Indeed, the majority of the control non-optically stimulated Kiss-Cre mice in estrus exhibited no LH pulse or intermittently 1 pulse during the 2.5 h sampling period (Fig. 2A&E). Similarly, no LH pulses or occasionally 1 LH pulse was observed in the 60 min control period in the optically stimulated mice (Fig. 2E; white bars). Optic stimulation at 0.5 Hz failed to induce LH pulses (Fig. 2B&E). In contrast, increasing the low-frequency continuous stimulation to 1 Hz evoked regular LH pulses (Fig. 2C&E, while 5 Hz resulted in a further, statistically significant (p < 0.05), increase in LH pulse frequency (Fig. 2D&E), all in line with our theoretical predictions (Fig. 1C). Continuous optogenetic stimulation (5 Hz) of AAV-injected wild-type C57BL/6 estrous mice (n = 3) failed to induce LH pulses (data not shown) further confirming that increasing the basal activity in the ARC KNDy neuronal population via low frequency continuous stimulation is sufficient to evoke and sustain LH pulsatile dynamics.

**Figure 2:**
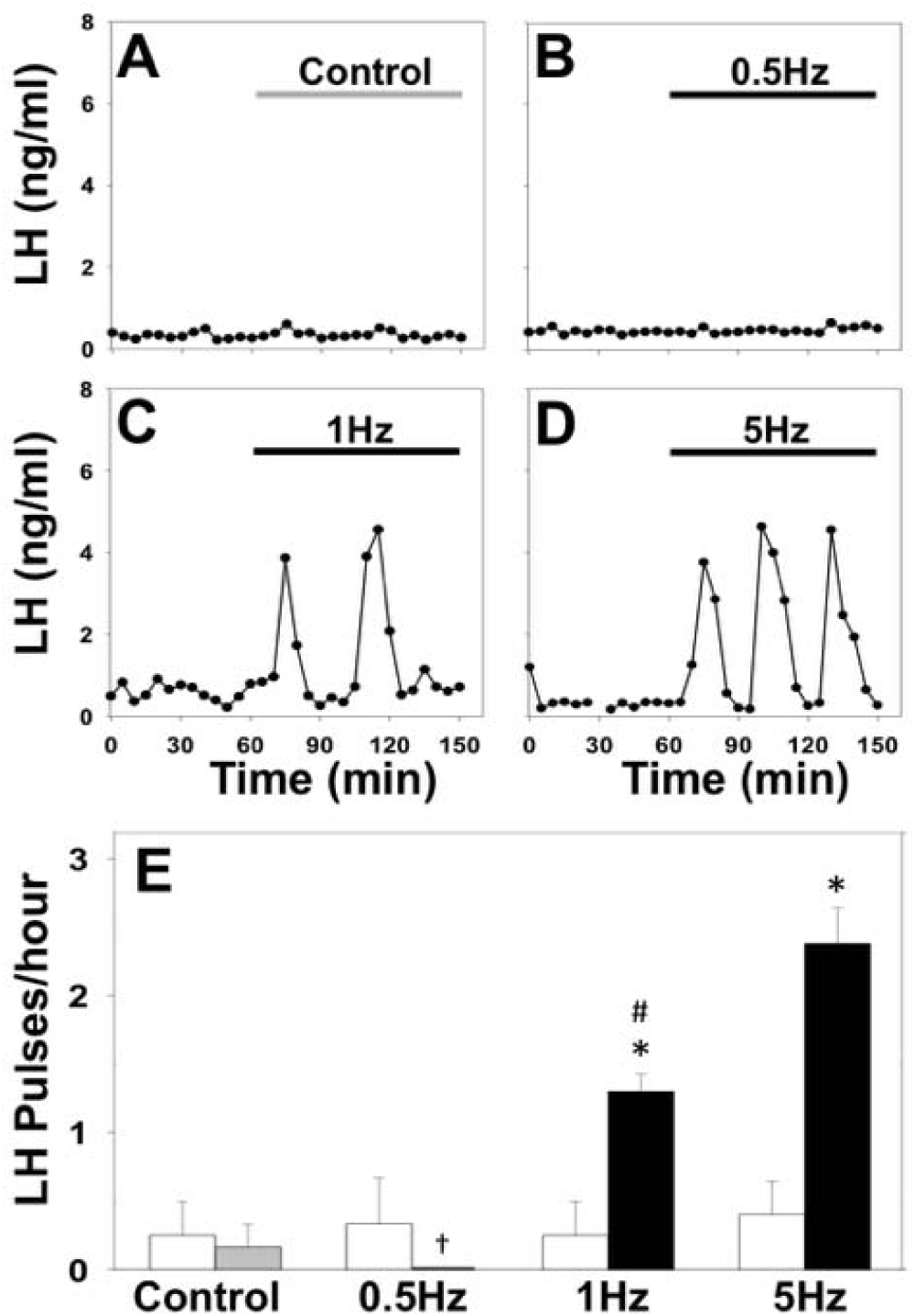
Optic stimulation of ARC kisspeptin neurons triggers LH pulses in estrous Kiss-Cre mice. (A-D) Representative examples showing LH secretion in response to no stimulation (grey bar) as control (A), or continuous blue light (473 nm, 5-ms pulse width, black bar) activation of kisspeptin neurons at 0.5 Hz (B), 1 Hz (C) or 5 Hz (D). (E) Summary showing for each group mean ± SEM LH pulse frequency over the 60min control period (white bars) and over the subsequent stimulation period (black bars). *P < 0.05 vs pre-stimulation; ^#^P < 0.05 vs stimulation at higher frequency. †Absence of LH pulses in response to 0.5 Hz stimulation. n = 4-6 per group.

### Levels of Dyn and NKB signaling control the response of the system to optic stimulation

The above theoretical and experimental results reveal a characteristic tipping-point behavior of the system, where a small increase in the basal activation levels is sufficient to trigger robust pulsatile dynamics (22). Our model predicts that such behavior emerges as a result of the non-linear positive and negative feedback interactions that are mediated through NKB and Dyn signaling respectively. Therefore to test the active role of NKB and Dyn signaling on pulse generation, we next combined optogenetic stimulation with neuropharmacological perturbations of the two pathways.

#### Disrupting Dyn signaling increases the sensitivity of the system to optic stimulation

Our model predicts that disruption of Dyn signaling should enable pulsatile dynamics over a wider range of optic stimulation frequencies (Fig. 3A). Intuitively, such disruption will reduce the strength of negative feedback in the system and consequently lower optic stimulation frequencies would suffice to excite the bi-stable neuronal population and set the relaxation oscillator in motion. To test this prediction *in-vivo* we repeated the optogenetic stimulation protocol at 0.5Hz together with nor-binaltorphimine (nor-BNI), a selective kappa opioid receptor (KOR) antagonist, to block Dyn signaling. Although 0.5Hz had previously failed to induce LH pulses (Fig. 4C&F), the addition of nor-BNI (bolus intra-cerebroventricular injection of 1.06 nmol over 5 min, followed by a continuous infusion of 1.28 nmol over 90 min) evokes a statistically significant increase in LH pulse frequency to approximately 1.6 pulses/hour (Fig. 3F). Intra-cerebroventricular injection of nor-BNI alone had no effect on LH pulse frequency (Fig. 3D&F).

**Figure 4:**
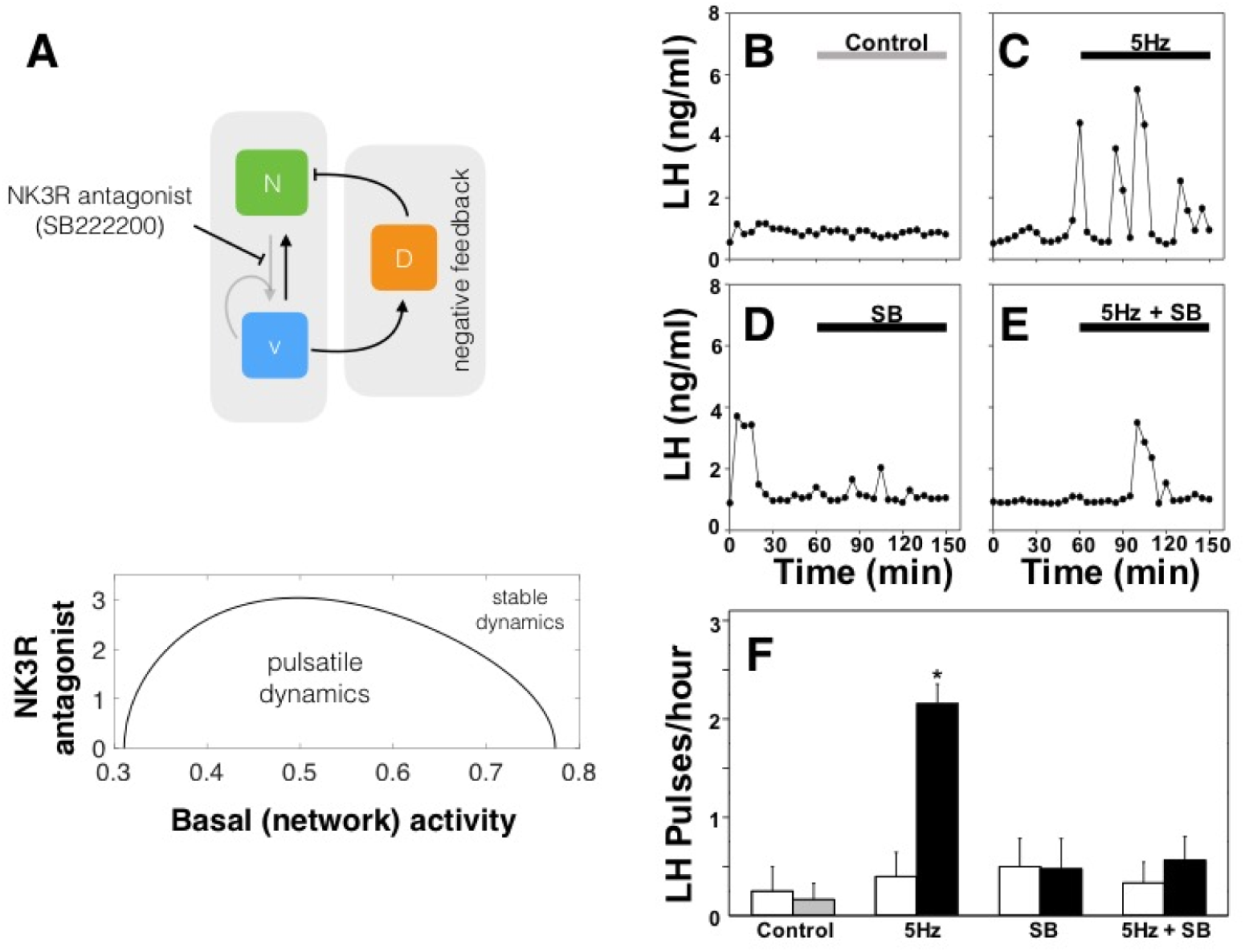
Disrupting NKB signaling desensitize the system to optic stimulation. (A) The model predicts that weakening bi-stability by blocking NKB signaling should restrict the range of optic stimulation frequencies that trigger pulsatility. SB222200 (Sb), an NKB receptor (TACR3) antagonist, was used to block NKB signaling *in-vivo*. (B-F) Representative examples showing LH secretion in response to no treatment (grey bar) as control (B), continuous blue light (473 nm, 5-ms pulse width, black bar) activation of kisspeptin neurons at 5 Hz (C), SB222200 treatment (bolus intra-cerebroventricular injection of 6 nmol over 5 min, followed by a continuous infusion of 9 nmol over 90 min) (D) combined SB222200 treatment and optic stimulation at 0.5Hz (E). (F) Summary plot showing for each group mean±SEM LH pulse frequency over the pre-treatment (white bars) and treatment period (black bars). *P < 0.05 vs pre-treatment. n = 4-6 per group.

**Figure 3:**
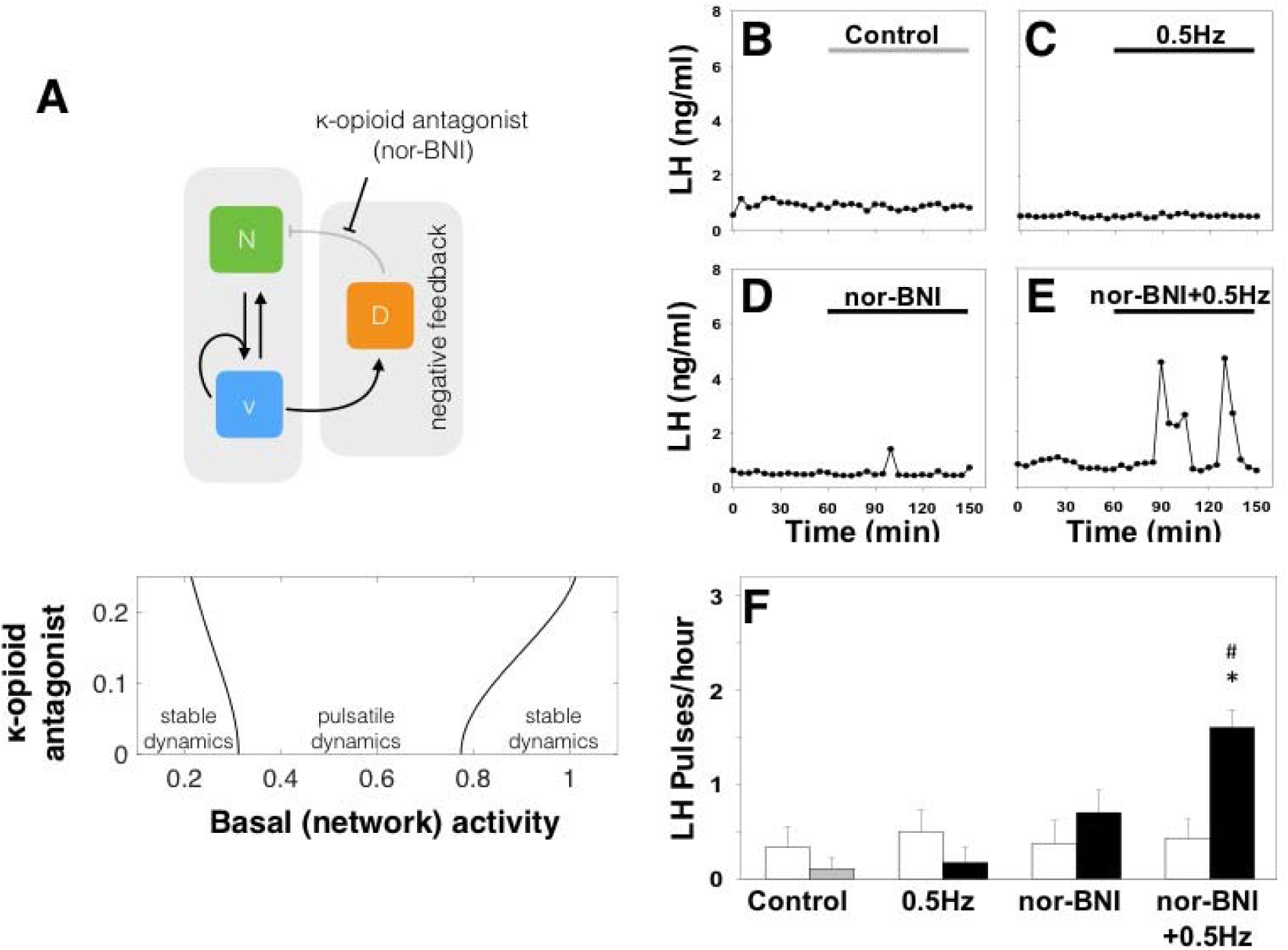
Disrupting Dyn signaling increases the sensitivity of the system to optic stimulation. (A) The model predicts that reducing the strength of negative feedback by partially blocking Dyn signaling should enable pulsatile dynamics over a wider range of optic stimulation frequencies. Nor-BNI, a Dyn receptor (kappa-opioid) antagonist, was used to block Dyn signaling *in-vivo*. (B-F) Representative examples showing LH secretion in response to no treatment (grey bar) as control (B), continuous blue light (473 nm, 5-ms pulse width, black bar) activation of kisspeptin neurons at 0.5 Hz (C), nor-BNI treatment (bolus intra-cerebroventricular injection of 1.06 nmol over 5 min, followed by a continuous infusion of 1.28 nmol over 90 min (D) combined nor-BNI treatment and optic stimulation at 0.5Hz (E). (F) Summary plot showing for each group mean±SEM LH pulse frequency over the pre-treatment (white bars) and treatment period (black bars). *P < 0.05 vs pretreatment; #P < 0.05 vs only optic stimulation treatment. n = 4-6 per group.

#### Disrupting NKB signaling desensitizes the system to optic stimulation

Our model predicts that disruption of NKB signaling should desensitize the system to external optic stimulation (see Fig. 4A). NKB signaling is key for pulsatile behavior as it enables positive feedback interactions within the population and therefore promotes bi-stability. Hence disrupting NKB signaling ought to decrease the propensity of the system to get excited into the pulsatile regime by external stimulation. To test this prediction *in-vivo* we repeated the optogenetic stimulation protocol at the highest frequency (5Hz), and used SB222200, a selective TACR3 antagonist, to block NKB signaling. Although 5Hz had previously induced a high frequency LH response (Fig. 4C&F), we observe that antagonist addition (bolus intra-cerebroventricular injection of 6 nmol over 5 min, followed by a *continuous* infusion of 9 nmol over 90 min) completely blocked the increased LH pulse frequency (Fig. 4E&F). Intra-cerebroventricular injection of SB222200 alone had no effect on LH pulse frequency (Fig. 4D&F).

## Discussion

Motivated by recent experimental evidence, we developed and studied a mathematical model of the KNDy neural population in the ARC, a population proposed to comprise the core of the GnRH pulse generator (23, 24). Our model demonstrates that the KNDy population can indeed produce and sustain pulsatile dynamics working as a relaxation oscillator. On the one hand, auto-stimulation via NKB signaling allows the population to behave as a bi-stable switch, firing either at a high or low rate. Moreover, basal neuronal activity and negative feedback through Dyn signaling allow the population to switch between the two activity states in a regular manner, giving rise to pulses of neuronal activity. Using global sensitivity analysis, we found that this mechanism of pulse generation is robust to parameter perturbations (SI, Fig. S3). In fact, co-variation of parameters governing, for example, the magnitude of basal activity, and the maximum secretion rates of NKB and Dyn is a more effective way of modulating the systems’ oscillatory behavior (amplitude and frequency). This multi-channel mode of efficient regulation is perhaps not surprising given the system’s crucial function, and hints that steroid feedback modulating the dynamics of the pulse generator over the reproductive cycle in female mammals is mediated through multiple, possibly interlinked, pathways.

Following model predictions, we explored the effect of continuous optogenetic activation of the KNDy population on LH pulse dynamics, a proxy for GnRH pulse dynamics. The model predicts that pulsatile dynamics of the system critically depend on the levels of basal activity in the KNDy population, and highlights the tipping-point behavior of the system as basal activity is modulated. For example, starting from a silent state of the system, introducing low levels of basal activity can reinstate pulses and increase their frequency. We have tested these model predictions *in-vivo* using optogenetics (24-26), and showed that we are able to directly control the generation and frequency of LH pulses in estrous mice by selectively exciting KNDy neurons in a continuous fashion at 1Hz and 5Hz. It is important to note that the model has informed the design of the experiments by suggesting the use of low-frequency optic stimulation for the first time in investigating LH pulse generation and in contrast to previous studies (24-26). Our results suggest that inhibitory or excitatory synaptic signaling within the KNDy neural population have a drastic effect on GnRH/LH pulse dynamics. We speculate that this enables KNDy neurons to integrate and transmit information regarding the overall state of the organism that is relevant for reproduction: for example, information on the emotional state and stress level through synaptic connections originating at the level of the amygdala (27), a key limbic brain structure; or information regarding the nutritional state of the organism through connections from Agouti-related peptide (AgRP)-expressing neurons in the hypothalamus (28).

The model predicted that the systems’ pulsatile behavior emerges as a result of the non-linear positive and negative feedback interactions that are mediated through NKB and Dyn signaling respectively. Using experimental protocol suggested by model analysis we showed that the response of the system to external optic stimulation indeed follows our NKB and Dyn signaling predictions. Our results highlight the need for a quantitative understanding of how the sex-steroid milieu affects the NKB and Dyn signaling pathways in the KNDy population. Such an understanding will lead to more accurate interpretation of results from *in-vivo* neuropharmacological perturbation experiments in various animal models and will shed light on the mechanisms underlying the regulation of pulsatile LH secretion in various natural settings such as lactational amenorrhoea or pharmaceutic interventions including the hormone contraceptive pill. We envision that as hormonal measurement techniques advance, enabling accurate, real-time readouts from individuals at low cost, such predictive mathematical models would be a valuable tool for understanding of reproductive physiopathology.

## Materials and methods

### Bifurcation analysis and numerical experiments

Bifurcation analysis of the coarse-grained model was performed using AUTO-07p (29). Both the full network model and coarse-grained model were simulated in Matlab using function ode45 (explicit Runge-Kutta (4, 5) solver).

### Animals

Breeding pairs of Kiss-Cre heterozygous transgenic mice (30) were obtained from the Department of Physiology, Development and Neuroscience, University of Cambridge, UK. Litters from the breeding pairs were genotyped by polymerase chain reaction (PCR) analysis. Adult female mice (8-14 wk old; 25-30g) heterozygous for the Kiss-Cre transgene or wild-type C57BL/6 littermates, with normal pubertal development and estrous cyclicity, were used. Mice were housed under a 12:12 h light/dark cycle (lights on 0700 h) at 22 ± 2 °C and provided with food (standard maintenance diet; Special Dietary Services, Wittam, UK) and water ad libitum. All animal procedures performed were approved by the Animal Welfare and Ethical Review Body (AWERB) Committee at King’s College London, and in accordance with the UK Home Office Regulations.

### Surgical procedures

Surgical procedures for stereotaxic injection of AAV9-EF1-dflox-hChR2-(H134R)-mCherry-WPRE-hGH (4.35 x 1013 GC/ml; Penn Vector Core) to express channelrhodopsin (ChR2) in ARC kisspeptin neurons were performed under aseptic conditions with general anesthesia induced by ketamine (Vetalar, 100 mg/kg, i.p.; Pfizer, Sandwich, UK) and xylazine (Rompun, 10 mg/kg, i.p.; Bayer, Leverkusen, Germany). Kiss-Cre female mice (n = 7) or wide-type (n = 3) were secured in a David Kopf Motorized stereotaxic frame and surgical procedures were performed using a Robot Stereotaxy system (Neurostar, Tubingen, Germany). A small hole was drilled into the skull at a location above the ARC. The stereotaxic injection coordinates used to target the ARC were obtained from the mouse brain atlas of Paxinos and Franklin (31) (0.3 mm lateral, 1.2 mm posterior to bregma and at a depth of 6.0 mm). Using a 2-μL Hamilton micro-syringe (Esslab, Essex, UK) attached to the Robot Stereotaxy, 1 μl of the AAV-construct was injected unilaterally into the ARC at a rate of 100 nl/min. The needle was left in position for a further 5 min and then removed slowly over 1 min. A fiber optic cannula (200 Hm, 0.39NA, 1.25mm ceramic ferrule; Thorlabs LTD, Ely, UK) was then inserted at the same coordinates as the injection site, but to a depth of 5.88 mm, so that the fiber optic cannula was situated immediately above the latter. Dental cement or a glue composite was then used to fix the cannula in place, and the skin incision closed with suture. A separate group of mice (n = 10) injected with the AAV construct, and fiber optic cannulae as described above, but additionally chronically implanted with an intra-cerebroventricular (icv) fluid cannulae (26 gauge; Plastics One, Roanoke, VA, USA) targeting the lateral ventricle (coordinates: 1.1 mm lateral, 1.0 mm posterior to bregma and at a depth of 3.0 mm), was used for the combined neuropharmacological and optogenetic studies. After surgery, mice were left for 4 weeks to achieve effective opsin expression. After a 1-wk recovery period, the mice were handles daily to acclimatize them to the tail-tip blood sampling procedure (32).

### Experimental design, and blood samplings for LH measurement

Prior to optogenetic stimulation, the very tip of the mouse’s tail was excised using a sterile scalpel for subsequent blood sample collection (21). The chronically implanted fiber optic cannula was then attached via a ceramic mating sleeve to a multimode fiber optic rotary joint patch cables (Thorlabs), allowing freedom of movement of the animal, for delivery of blue light (473 nm wavelength) using a Grass SD9B stimulator controlled DPSS laser (Laserglow Technologies, Toronto, Canada). Laser intensity at the tip of the fiber optic patch cable was 5 mW. After 1 h acclimatization, blood samples (5μl) were collected every 5 min for 2.5 h. After 1 h controlled blood sampling, continuous optic stimulation (5-ms pulse width) was initiated at 0.5, 1 or 5 Hz for 90 min. Controls received no optic stimulation. Kiss-Cre mice received the stimulation protocols in random order. Wild-type received 5 Hz optic stimulation only.

For the neuropharmacological manipulation of Dyn or NKB signaling with or without simultaneous optogenetic stimulation the animals were appropriately prepared as described above, but in addition an icv injection cannula with extension tubing, preloaded with drug solution (nor-BNI or SB222200 dissolved in artificial cerebrospinal fluid), was inserted into the guide cannula immediately after connection of the fiber optic cannula. The tubing was extended outside the cage and connected to a 10 μl syringe (Hamilton) mounted in an automated Harvard pump (Harvard Apparatus, Holliston, MA, USA) to allow remote microinfusion without disturbing the mice during the experiment. Five min before optic stimulation, icv administration of drug treatment commenced as a bolus injection over 5 min, followed by a continuous infusion for the remainder of the experiment. In the absence of optic stimulation the same icv regimen was used. The blood samples were processed by ELISA as reported previously (32). Mouse LH standard and antibody were purchased from Harbour-UCLA, USA, and secondary antibody (NA934) was from VWR International, UK. The intra-assay and inter-assay variations were 4.6% and 10.2%, respectively.

### Validation of AAV injection site

After completion of experiments, mice were anaesthetized with a lethal dose of ketamine and transcardially perfused with heparinized saline for 5 min, followed by 10 min of ice-cold 4% paraformaldehyde (PFA) in phosphate buffer (pH 7.4) for 15 min using a pump (Minipuls, Gilson, Villiers Le Bel, France). Brains were rapidly collected and postfixed sequentially at 4 °C in 15% sucrose in 4% PFA and in 30% sucrose in phosphate-buffered saline until they sank. Afterwards, brains were snap-frozen on dry ice and stored at −80 °C until processing. Brains were coronally sectioned (40-μm) using a cryostat (Bright Instrument Co., Luton, UK) and every third section was collected between −1.34 mm to −2.70 mm from the bregma. Sections were mounted on microscope slides, air-dried and cover slipped with ProLong Antifade mounting medium (Molecular Probes, Inc. OR, USA). The injection site was verified and evaluated using Axioskop 2 Plus microscope equipped with axiovision 4.7 (see also SI, Fig. S1). One of 17 Kiss-Cre mice failed to show mCherry fluorescence in the ARC and was excluded from the analysis.

### LH Pulses and Statistical Analysis

Detection of LH pulses was established by use of the Dynpeak algorithm (33). The effect of optogenetic stimulation on parameters of LH secretion was calculated by comparing the mean number of LH pulse per hour, within the 90 min stimulation/drug delivery period with the 60 min prestimulation/drug delivery control period. For the non-stimulated control animals, the same timepoints were compared. The mean number of LH pulse per hour, within the 90 min stimulation period, or equivalent, was also compared between experimental groups. Statistical significance was testes using one-way ANOVA followed by Dunnett’s test. P < 0.05 was considered statistically significant. Data are presented as the mean ± SEM.

## Supporting information

**S1 text.**

**Supporting Information.**

## Acknowledgments

KTA and MV gratefully acknowledge the financial support of the EPSRC via grant EP/N014391/1. KOB and SLL gratefully acknowledge the financial support of the MRC via grant MR/N022637/1.

